# Anaerobic degradation of acid red 73 by obligately aerobic *Aspergillus tabacinus* LZ-M though a self-redox mechanism

**DOI:** 10.1101/2022.05.16.492227

**Authors:** Xuan Yu, Simin Zong, Chunlan Mao, Aman Khan, Wenxue Wang, Hui Yun, Peng Zhang, Xiangkai Li

## Abstract

Fungi are potential biological resources for refractory organics degradation, but their anaerobic degradation of azo dyes are rarely reported. In this study, a fungus *Aspergillus tabacinus* LZ-M was isolated grown aerobically and degraded acid red 73 (AR73) with a decolorization rate of 90.28% in 5 days at 400 mg/L of concentration anaerobically. Metabolic pathway showed that AR73 was reduced into 2-hydroxynaphthalene and aniline then mineralized into CO_2_. The anaerobic self-redox process revealed electrons generated in carbon oxidation and transferred to -C-N= and -N=N, resulting in complete mineralization of AR73 in strain LZ-M. Data of transcriptome analysis showed that the benzene compounds produced from AR73 by declorizing reductase entered the catechol pathway and glycolysis process to mineralize. Enzymes involved in aromatics degradation, glycolysis processes, cytochrome C and quinone oxidoreductases were up-regulate, but the key reductase responsible to cleave AR73 to phenylhydrazine was not found. A novel enzyme Ord95 containing a glutamate S-transferase domain was identified in the unknown genes as a reductase which cleaving -C-N= in AR73 using NADH as electron donor, and three arginines key active sites. These observations reveal a new degradation mechanism of AR73 in strain LZ-M which would be potential candidate for treatment of azo dyes wastewater.

**Graphical abstract:** 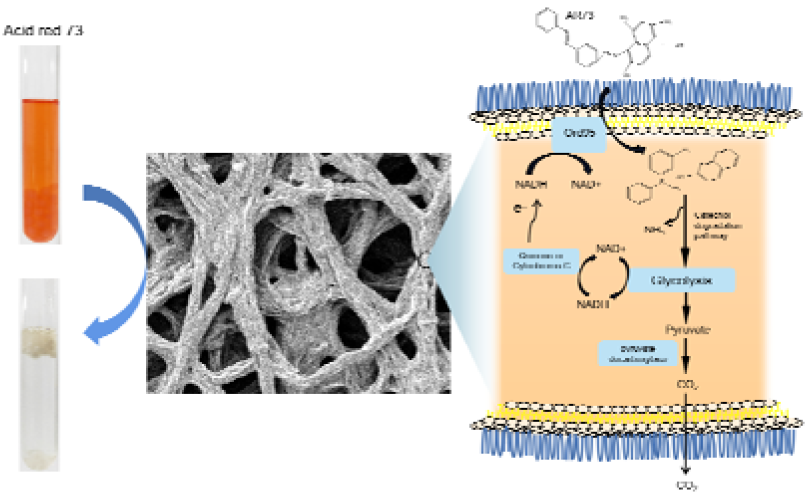

## 1. Introduction

Azo dyes are widely used as colorants in different fields of its properties, such as vivid colors, tinctorial strength, and ease of manufacturing, and approximately 450 000 tons of azo dyes are consumed annually (*1, 2*). Among 15% of dyes used in the industries are discharged into wastewater (*3*). Dyes are refractory, toxic and mutagenic, inevitably causing extensive pollution in both aquatic and terrestrial systems (*4*). In recent decades, biotechnologies have shown efficiency and affordability advantages in treatment of wastewater containing toxic and refractory dyes (*5*). Especially, bacteria-mediated treatments have emerged to eliminate azo dyes from wastewater (*6, 7*). Nevertheless, the bacterial treatment can only decolorize, but not able to achieve the complete mineralization of azo dyes (*3, 8*). The symmetrically breakdown of azo bond by bacteria is easily to accumulate aromatic amines, which are toxic to organisms and are stubborn for biodegradation (*9*). The asymmetric breakage of azo bond or complete mineralization of azo dyes by microorganisms are rare. Therefore, it is essential to find the microorganisms that can degrade the azo dyes to non-toxic substances.

Fungi, as a decomposer showed significant capabilities in the degradation of toxic and refractory organics including lignin, cellulose, pharmaceuticals, polycyclic aromatic hydrocarbons, and etc. (*10, 11*). Fungi secret non-specific oxidases and completely mineralize these pollutants into CO_2_ (*12*). For example, benzo[a]pyrene is mineralized to CO_2_ by the white rot fungus *Pleurotus ostreatus* (*13*). *Aspergillus versicolor* LH1 completely degrade methyl red at 200 mg/L of concentration (*2*), and *A. niger* decolorize acid blue 161 up to 58% (*3*). However, the decolorization of azo dyes are commonly more effective under anaerobic than aerobic conditions (*9*). This is because of the azo reduction enzyme for the initial cleavage of the azo bond sensitive to oxygen (*2, 14*). Similarly, the electron withdrawing nature of the azo bond impedes the susceptibility of dye molecules to oxidative reaction and, thus, azo dyes show resistance to aerobic biodegradation (*14*). Fungi commonly grow under aerobic condition, and degradation of azo dyes under anaerobic condition was less reported.

Some reports suggest that aerobic microorganisms have potential for anaerobic reaction (*15*). For example, Lysinibacillus sp. NP05 was capable of degrading polychlorinated biphenyls under anaerobic condition (*15*). Pseudomonas denitrificans G1 grew under the aerobic condition, and could achieve effective denitrification under the anaerobic conditions (*16*). An obligately aerobic bacterium, Zetaproteobacteria, has an anaerobic metabolic pathway by genome analysis (*17*). This indicates that enzymes associated with anaerobic function are secreted by this aerobic microorganism. There are evidences that auxiliary anaerobic metabolism exists in obligately aerobic fungi (*18*). An obligately aerobic fungus, A. nidulans, could survive under anaerobic conditions and exists the alcoholic fermentation pathway (*18, 19*). So, fungi have the potential to degrade azo dyes under anaerobic conditions.

The bio-degradation process of organic matter is usually accompanied with oxidation and reduction reactions. For example, lignin, plastic and dyes are oxidatively depolymerized by peroxidase (*20*). Anaerobic denitrification of organic nitrogen is a reduction process (*21*). Some organic matters are degraded synchronously by oxidation and reduction in a self-redox process. For instance, bioconversion of organic matter to methane/H_2_ during anaerobic biological fermentation (*22*). In a fermentation process, electrons were generated during butyrate oxidation and transferred to H^+^ to generate acetate and H_2_ (*23*). In this study, a fungus identified as *Aspergillus tabacinus* LZ-M could grow aerobically and degrade acid red 73 (AR73) anaerobically. The analysis of metabolic pathway illustrated that LZ-M had a strong ability to mineralize AR73 via self-redox. A novel decolorizing reductase was excavated and its degrading mechanism was revealed. This study provides a new microbial resource for the biodegradation of azo dyes.

## 2. Materials and methods

### 2.1 Strain isolation and culture

To isolate microorganisms adapted to low-nutrient wastewater, a piece of parafilm (PM996) (1 cm × 1 cm) was floated in 100 mL of carbon-free liquid medium (1 g/L NaCl, 1 g/L K_2_HPO_4_, 1 g/L KH_2_PO_4_) and inoculated with 1 mL soil sample that was collected in the arboretum of Gansu Academy of Membrane Science and Technology in Gansu Province, China. After stationary culture for 30 days at room temperature (25-28 °C), flocculent microorganisms were growing around the parafilm. The flocculent microorganisms were transferred into the solid medium of Potato Dextrose Broth Medium (PDB, Solarbio, China), culture for 3 days at 30 °C. Pick spores into the solid medium of PDB for culturing until pure colonies were formed. For spore collection, using 10 mL 25% glycerol solution to elute the spores on the surface of the colonies in a solid culture plate, and sucking the spore liquid into the cryogenic tubes and stored it at −80 °C.

The aerobic growth of this strain was conducted in 250-mL conical flasks containing 150 mL of liquid PDB medium with inoculating 150 uL of spore suspension, and mycelia pellets were formed after culturing for 2.5 days at 30 °C and 200 rpm. The spore concentration is about 10^4 /mL, calculated as the number of mycelia pellets formed by liquid culture using serial dilution. For activated culture, the amount of spore inoculum is 10.

### 2.2 Chemicals

Acid red 73 was purchased from Sigma-Aldrich (St. Louis, USA). Methanol and ethyl acetate were HPLC grade. Other reagents were purchased from Solarbio Science & Technology Co.,Ltd. Beijing, China.

### 2.3 Anaerobic decolorization of fungi

Mycelia pellets from 300 mL aerobic activated medium were collected by filtration and washed for 5 times using ddH_2_O and then used for anaerobic degradation experiment. One liter of basal mediun (BM) contained 1 g/L NaCl, 1 g/L KH_2_PO_4_, 1 g/L K_2_HPO_4_, and pH 7 was used to suspend the mycelium pellets. To analyze the ability of fungi to decolorize AR73 under anaerobic condition, AR73 solutions were added into the medium to arrive the concentration of 50 mg/L. The 100 mL of the mixed mycelium suspension was added to 250 mL anaerobic flasks respectively. Anaerobic flasks was filled with nitrogen for 20 min and stationary incubated in an anaerobic incubator at 30 °C (n=3). For anoxic culture, anaerobic flasks without nitrogen pouring were incubated at 30 °C in static phase. For aerobic culture, flasks were rotationally incubated at 30 °C and 200 rpm. Experiment used sterile medium as controls. The samples were taken at different time interval for analysis.

### 2.4 Effect of operation parameters on biodecolorization

The biodecolorization performance of AR73 by strain LZ-4 was evaluated under different culture conditions. Mycelia pellets from 900 mL aerobic activated medium were suspended in 3 L BM and divided into anaerobic bottles with 100 mL per bottle. To examine the effect of additional carbon sources, 1 g/L of soluble starch, glucose, potato, tyrptone, PDB medium were added into BM containing 50 mg/L AR73, respectively. To examine the effect of pH, the initial pH was adjusted to 3, 4, 5, 6, 7, 8, 9, 10, and 10.5. In AR73 load experiments, the initial AR73 concentrations ranged from approximately 50-500 mg/L. To evaluate the ability of the strain LZ-M for other dyes and refractory organics removal, 50 mg/L methyl orange, 50 mg/L neutral red, 30 mg/L malachite green, 10 mg/L metronidazole, 30 mg/L furazolidone and 30 mg/L 3,5-dinitrosalicylic acid were added into BM, respectively. In order to test the long-term anaerobic decolorization by strain LZ-M, 2.5 mL of 2g/L AR73 was added to the anaerobic bottle when the color of the dye in the culture solution was disappeared, continuously, until occurring incomplete depigmentation after 24 hours. Unless the single-factor was adjusted as per the experimental design, the initial concentration of AR73 concentration was 50 mg/L. The fungal suspension was incubated for 5 days at 30 °C under anaerobic condition, and samples were collected to determine the concentrations of substrate. All experiments were performed in three duplicate.

### 2.5 LC/MS analyses of metabolites

After anaerobic degradation of 50 mg/L AR73 by strain LZ-M for 3 days, the culture supernatant was collected and sent to Wuhan Metware Biotechnology Co.,Ltd. for LC/MS analysis. Metabolic identification information was obtained by searching the company’s database.

### 2.6 Analysis of ITS sequencing and transcriptome sequencing

Mycelia pellets of strain LZ-M after aerobic growth were collected and send to Shanghai Majorbio Bio-pharm Technology Co.,Ltd (Shanghai, China) for DNA extraction and ITS sequencing.

To prepare the sample for transcriptome analysis, mycelia pellets of *A. tabacinus* LZ-M after aerobic growth were transferred to BM containing 50 mg/L AR73, filled with nitrogen for 20 min and cultured for 3.5 hours under anaerobic condition at 30 °C. The culture cells were cultured at 200 rpm under aerobic condition were used as a control. After that, mycelia pellets were collected and send to Shanghai Majorbio Bio-pharm Technology Co.,Ltd (Shanghai, China) for RNA extraction and transcriptome sequencing.

### 2.7 Cloning of unknown genes

The genes sequence of Ord95 and Ord118 was represented in supplementary and were clone as follows; the Ord95 fragment was amplified from the transcriptome RNA by polymerase chain reaction (PCR) using the primers Ord95F: 5’-CCGGAATTCATGTCAGATTCCACGCTCTACC-3’ and Ord95R: 5’-CCGCTCGAGGCCCTCCAACGCATCTTC-3’. The Ord118 fragment was PCR using the primers Ord118F: 5’-CCGGAATTCATGGCTACTCAAGCTATTCAC-3’ and Ord118R: 5’-CCCAAGCTTGCGGTGATGCAGCATGTC-3’. The Ord95 and Ord118 fragments were digested with restriction endonucleases and inserted into plasmid pET-28a, respectively. The recombinant plasmids pET28/Ord95 and pET28/Ord118 were transformed into *E.coli* Rosseta (DE3). For protein-induced expression, the cells were grown in Luria-Bertani medium containing Kanamycin (50 μg/mL) and Chloramphenicol (30 μg/mL) at 37 °C. When cells grown to an OD_600_ of 0.6, 0.2 mM isopropyl β-D-1-thiogalactopyranoside was added to the medium and was then cultured for 20 h at 16 °C. Purification of protein followed as a previous study (*24*). The purified protein was detected using SDS-PAGE, and protein concentration was detected using Pierce BCA Protein Assay Kit (Thermo Fisher Scientific, United States).

Various Ord95 mutants were constructed by Phusion™ Site-Directed Mutagenesis Kit (Thermo Fisher Scientific, United States). The procedures for the expression and purification of these variants were similar to that of wild type Ord95.

### 2.8 Enzyme assay of Ord118, Ord95 and its mutants

To explore NADH dehydrogenase activity of purified Ord118 and Ord95, the enzyme assay was performed in 3 mL 20 mM Tris-HCl buffer (pH 7.0), containing 0.28 mg/L protein, 200 μM NADH and was kept at 37 °C for 120 min. The reaction solution was placed in a cuvette, and the absorbance change of NADH was detected by UV-Spectrophotometry (Hitachi U-3900H) at 340 nm in real time. To explore the AR73 declorizing activity of purified proteins, the enzyme assay was performed in 5 mL of 20 mM Tris-HCl buffer (pH 7.0) containing 0.28 mg/L protein, 25 mg/L AR73 and 200 uM NADH. The reaction solution was injected into vacuum tubes in an anaerobic incubator and kept at 37 °C for 48 h, and 200 uL reaction solution was taken out from the vacuum tube every 12 h to measure the absorbance of AR73 at 510 nm. Reaction with predenaturation protein treated for 15 min at 95 °C were used as controls.

### 2.9 Analytical methods

The surface structure of mycelia pellets of *A. tabacinus* LZ-M was observed by scanning electron microscopy (JEOL JSM-IT500LA). Spectrophotometry was used to measure the concentrations of AR73, methyl orange, neutral red and malachite green at wavelength of 510 nm, 462 nm, 523 nm and 620 nm, respectively. Concentrations of metronidazole, furazolidone, and 3, 5-dinitrosalicylic acid were determined using UV-Spectrophotometry (Hitachi U-3900H) at wavelength of 320 nm, 370 nm and 350 nm, respectively.

Phenylhydrazine and aniline in the culture and enzymatic reaction solutions were detected by high-performance liquid chromatography (HPLC, Agilent 1260). HPLC analysis utilized an Agilent Eclipse XDB-C18 (4.6 mm ×250 mm, 5μm), the mobile phase was methanol/H_2_O solution (40:60, v/v) at a flow rate of 1.5 mL/min. An injection volume of 10 uL was used and the UV detection was performed at 280 nm. The consent of CO_2_ produced in headspace of anaerobic bottle was detected by gas chromatography (Agilent 6890N). The injector was set at 240 °C, the amount of gas sample in each injection was 20 uL, and a split less mode (60 mL/min) was used.

### 2.10 Statistical analysis

Statistical analyses were performed using SPSS v.17 software and Excel 2010. One-way Analysis of Variance was used to assess differences in the abundance of taxa and then values are presented as the mean ± standard error. Subsequently, Tukey’s test was used to determine the sample means that were significantly different.

## 3. Results and discussion

### 3.1 The identification of anaerobic AR73 degradation fungus

A fungus growing on parafilm surface in liquid medium was isolated which has 100% similarity to *Asperillus tabacinus* NRRL 4791, and named as *Aspergillus tabacinus* LZ-M (Fig. 1A). The isolated strain LZ-M was grow at 150-250 rpm, where the biomass growth rate increased with the rotation speed (Fig. S1), while less than 100 rpm strain LZ-M growth was inhibited. This result indicated that strain LZ-M was obligately aerobic fungus. The mycelium pellets of strain LZ-M was formed after aerobic growth, and placed it in an anaerobic environment. After 24 h, bubbles were generated in the culture medium, carrying the mycelium to the top of liquid (Fig. 1B). Furthermore, the dye AR73 was completely decolorized after 48 hours of incubation in anaerobic condition. Scanning electron microscope images confirmed the softening of hyphae caused by gas secretion (Fig. 1C). This data indicates that the aerobic strain LZ-M uses AR73 as a substrate for respiration and metabolism under anaerobic environment.

**Fig. 1.**
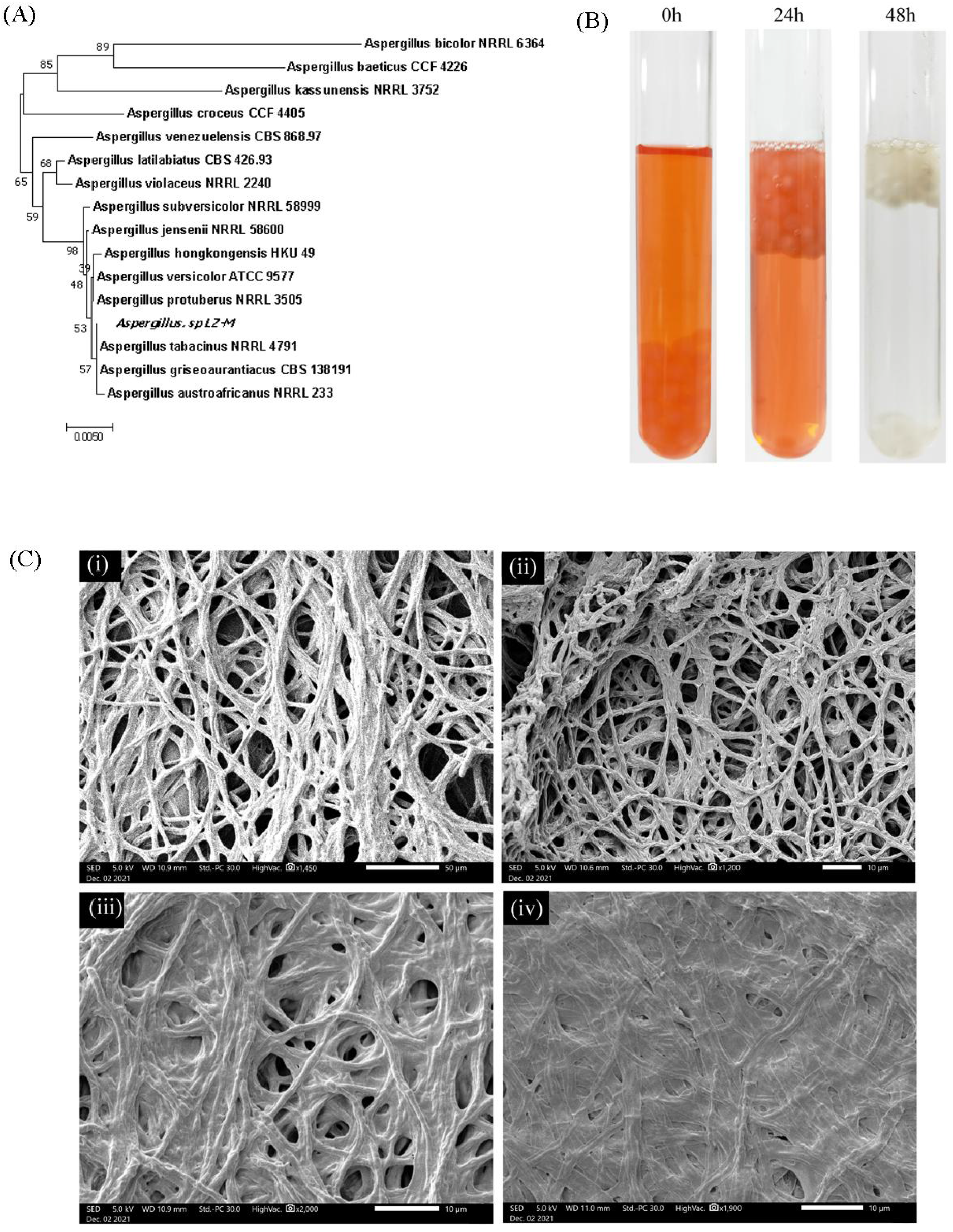
The identification of fungi LZ-M and its decolorization of acid red 73 under anaerobic condition. (A) Phylogenetic tree of strain LZ-M based on ITS sequencing; (B) Anaerobic decolorization process of fungi LZ-M mycelium in anaerobic tubes; (C) Scanning electron microscope images of the surface structure of mycelium: (i) mycelium after aerobic culture for 48h; (ii) mycelium at initial time of anaerobic culture; (iii) mycelium after anaerobic culture for 24h; (iv) mycelium after anaerobic culture for 48 h.

*Aspergillus* fungi growing under aerobic condition (*2, 25*), which was consistent with *A. tabacinus* LZ-M reported in this study. The mycelium pellets and morphology are also similar to the *Aspergillus* species, such as *A. niger* and *A. versicolor* (*26, 27*). The mycelium of strain LZ-M produced gas and decolorized AR73, surviving in anaerobic environment. Similarly, anaerobic degradation of azo dyes by bacteria were also reported previously. For example, *S. oneidensis* MR-1 showed a decolorization capability for Cationic Red and *Pseudomonas sp*. SUK1 could decolorize dye Red BLI (*6, 28*). This phenomenon suggests that *A. tabacinus* LZ-M can be used as a potential strain for degradation of the azo dyes as similar as bacterial species.

### 3.2 The characteristics of dye degradation by *A. tabacinus* LZ-M

The decolorization ability of *A. tabacinus* LZ-M was compared under aerobic and anaerobic conditions (Fig. 2A). The results showed that strain LZ-M completely decolorized (96.8%) of AR73 in 50 mg/L of concentration at 40 h under anaerobic conditions. However, under aerobic conditions, mycelium does not have the ability to decolorize. The effect factors on anaerobic decolorization efficiency were analyzed. The addition of organic carbon sources, including glucose, PDB medium, tryptone, potato and soluble starch can promote strain LZ-M for decolorization. Among them, soluble starch increased the decolorization ratio from 36.21% to 93.8% within 18 h of incubation (Fig. 2B). The different initial concentrations of AR73 were tested (Fig. 2C). Strain LZ-M completely degraded AR73 in 200 mg/L while the removal rates were decreases to 96.33, 90.28 and 58.72 % in the 300, 400, and 500 mg/L of concentrations respectively. The anaerobic reaction achieved complete decolorization of 50 mg/L AR73 at a wide pH range of 4-10.5 and the removal rate was decreased to 43.09% in pH 3 (Fig. 2D). In the anaerobic degradation of multi-batch, this strain could completely degrade 50 mg/L AR73 for 5 times, and the decolorization efficiency began to decline at 5th addition (Fig. 2E). Degradation ability test of strain LZ-M on other organic matter under anaerobic condition was shown in Fig. 2F. The results revealed that strain LZ-M degraded 99.98%, 75.87%, 42.55%, 50.315%, 87.375% and 57.43%, of 50 mg/L methyl orange, 50 mg/L neutral red, 30 mg/L malachite green, 10 mg/L metronidazole, 30 mg/L furazolidone, and 30 mg/L 3,5-dinitrosalicylic acid within 5 days respectively. The content of these organics in aerobic culture was not changed (data not shown).

**Fig. 2.**
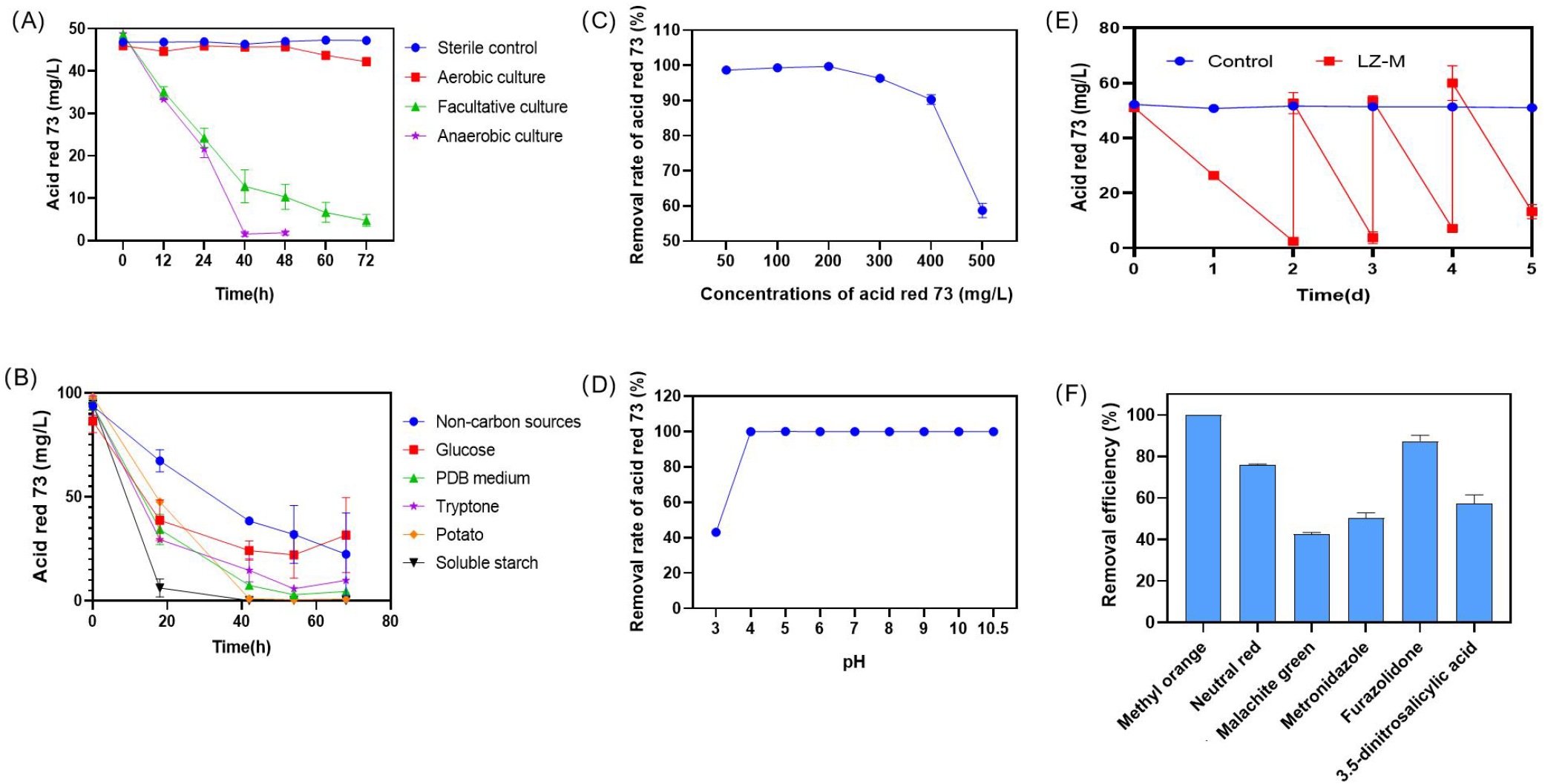
Effect factors of anaerobic decolorization of AR73 by *A. tabacinus* LZ-M and its degradation ability on other organic matter. (A) Variation curve of AR73 under aerobic and anaerobic conditions at different time; The effect of different carbon sources (B), AR73 concentrations (C) and pH (D) on decolorization; (E) Continuous decolorization of AR73 with multiple addition; (F) Anaerobic degradation of *A. tabacinus* LZ-M on different organic matters. All reaction experiments were

White rot fungus *Schizophyllum* adsorbs Acid Red 18 (100 mg/L) with a decolorization rate of 27% within 120 h (*29*). Another fungi *A. oryzae* showed 72.38% biosorption decolorization of Acid Red 337 (200 mg/L) (*30*). In contrast, this research reported the complete decolorization of 100-200 mg/L AR73 by strain LZ-M, showing a higher decolorization efficiency compared to the absorption of *Schizophyllum* and *A. oryzae*. The decolorization of strain LZ-M was also compared with the bacteria strains. *Stenotrophomonas sp*. BHUSSp X2 can achieve 90% decolorization of 500 mg/L Acid Red 1, while the effective decolorization is limited to pH 7-8 (*31*). *Bacillus thuringiensis* can achieve 60% decolorization of 500 mg/L acid red 119 (*32*). In this study, the strain LZ-M decolorized AR73 of 500 mg/L with a removal rate of 58.72% at 4-10.5 of pH ranges. The capability of stain LZ-M for decolorizing high concentrations of acid red under anaerobic condition was comparable to that of *Bacillus thuringiensis* (*32*), and the pH range of stain LZ-M for decolorizing was wider than *Stenotrophomonas sp*. BHUSSp X2 (*31*). These results showed the application advantage of *A. tabacinus* LZ-M in the environmental biotechnology, such as decolorizing industrial dyes with high concentrations and changing pH. This continuous multi-batch decolorization experiment demonstrated the potential of strain LZ-M in dye wastewater treatment. And its degradation on a broad of organics indicated that it is a promising candidate in pollutant treatment.

### 3.3 The degradation pathway of AR73 in *A. tabacinus* LZ-M

In order to analyze the anaerobic degradation pathway of AR73, the degradation products of AR73 by *A. tabacinus* LZ-M were detected by LC-MS. The results showed that 17 compounds were identified in the AR73 degradation (Table S1). Putative degradation pathway of AR73 was shown in Fig. 3A. The first and second step of degradation of AR73 was to produce phenylhydrazine and hydroxynaphthalene by the cleavage of -C-N= bonds through hydrogenation reduction. And then, phenylhydrazine was deaminated to aniline. Hydroxynaphthalene and aniline was hydroxylating and opening the benzene ring to generate carboxylic acid compounds. Carboxylic acid compounds were decarboxylated to produce CO_2_. The concentrations of phenylhydrazine and aniline was detected in the culture medium (Fig. 3B and C). With the increase of time, the concentration of aniline kept increasing, while the concentration of phenylhydrazine decreased after 48h, indicating that the intermediate metabolites phenylhydrazine was further denitrogenated into aniline. Gas produced in anaerobic reaction flasks was identified as CO_2_ by gas chromatography. The content of CO_2_ increased to 19.56% after 72 h in AR73 medium, and it was only 3.5% in the carbon-free control (Fig. 3D). This indicated that AR73 was mineralized into CO_2_ by *A. tabacinus* LZ-M under the anaerobic condition.

**Fig. 3.**
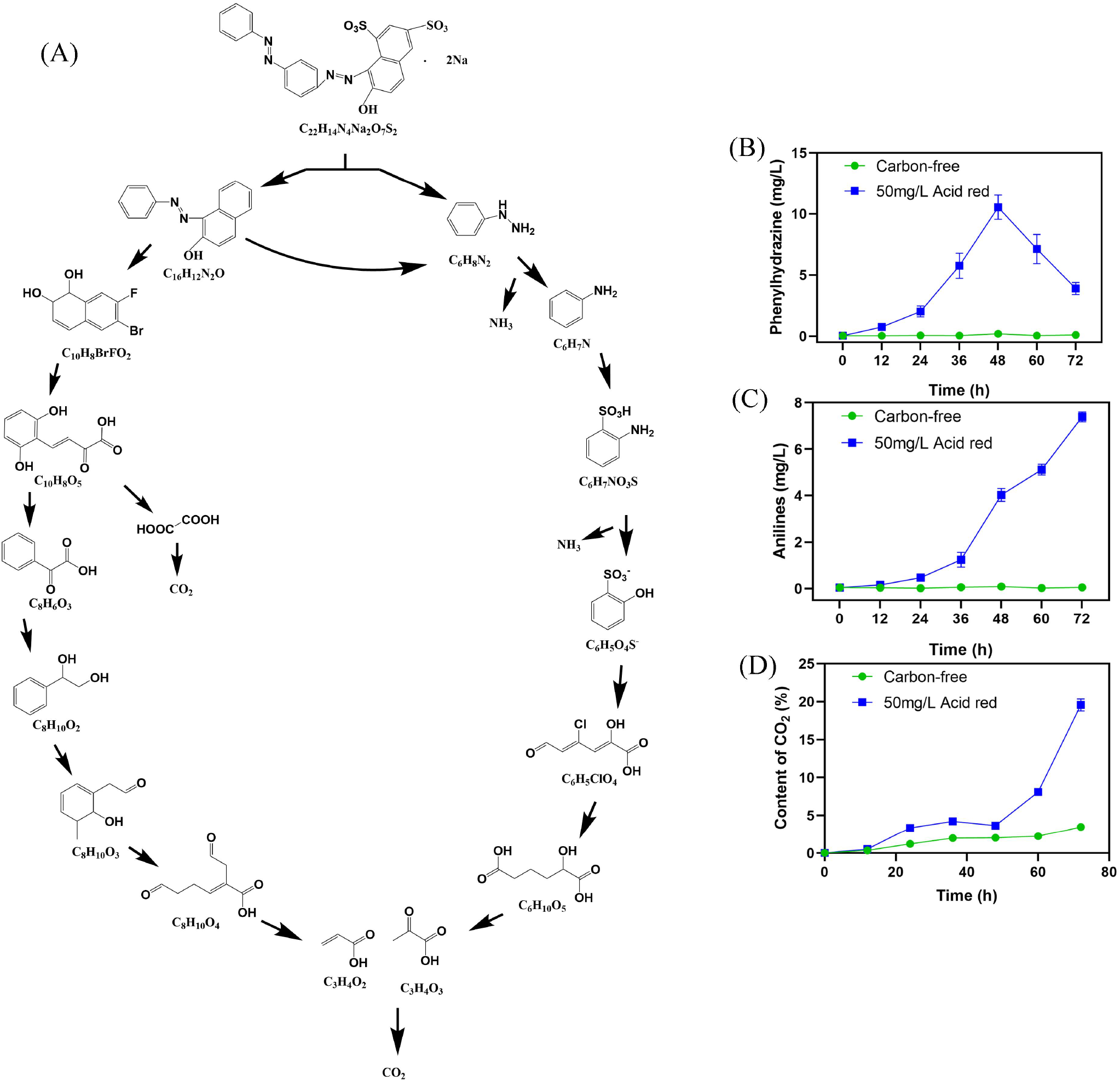
The degradation pathway of AR73 by *A. tabacinus* LZ-M. (A) Presumptive metabolic pathway of AR73 based on the products in LC-MS analysis results; The contents of phenylhydrazine (B), aniline (C) and CO_2_ (D) in anaerobic culture at differe

The degradation of AR73 to benzenes were mainly achieved by the cleavage of -C-N= bonds. *A. versicolor* LH1 degrades methyl red by breaking the -C-N= bond linked with benzene (*2*). Degradations of disperse yellow 3 and acid orange 7 by *Phanerochaete chrysosporium* are also achieved by the -C-N= cleavage (*33*). In this study, the -C-N= cleavage by the strain LZ-M is similar to these fungi. In reports about the bacterial strains, the degradation of Acid Red 1 by *Stenotrophomonas sp*. BHUSSp X2 is achieved by -N=N- cleavage (*31*). Azoreductase derived from *Sphingomonas xenophaga* BN6 degrades azo dyes also through -N=N- cleavage (*33*). This indicates that the degradation of azo dyes by strain LZ-M is different from the anaerobic degradation of these bacteria. In a previous study, naphthalene or benzene is hydroxylated and then benzene ring is opening at the hydroxyl position (*34*). This is consistent with the degradation pathway of hydroxynaphthalene and aniline in this study. The mineralization of benzene compounds to generate CO_2_ was achieved in white rot fungi and soil microbe (*35, 36*). The produce of CO_2_ indicated that the intermediate produces were mineralized to CO_2_. In totally, the degradation of AR73 is mainly achieved by deamination and decarboxylation. AR73 was the sole source of carbon and nitrogen, which acted as both an oxidant and a reductant in the reaction, so the degradation was achieved by self-redox process. The similar phenomenon was found in a previous study that *Syntrophomonas wolfei* can degrade butyrate to CO_2_ and H_2_ (*23*). During this self-redox reaction, the oxidant of AR73, and the reductant is the benzene compounds produced by AR73 decomposition.

### 3.4 Transcriptome analysis of fungi under aerobic and anaerobic conditions

In order to analyze the anaerobic degradation mechanism of *A. tabacinus* LZ-M, the differences of gene expression between aerobic and anaerobic conditions were analyzed by transcriptome sequencing. Under anaerobic conditions, a total of 4472 differentially expressed transcripts were found, including 2156 up-regulated transcripts and 2316 down-regulated transcripts (Fig. S2). The function types of up-regulated transcripts were shown in Table S2, and 1271 transcripts displayed unknown function, accounted for 51.81% of up-regulated transcripts. In addition, carbohydrate transport and metabolism including 167 transcripts was the most varied functional type. This indicated that carbohydrate transport and metabolism were significantly changed under anaerobic condition. Genes related to cytochrome C oxidoreductase, quinone oxidoreductase and the biodegradation of aromatics were significantly up-regulated (Fig. 4). A complete set of genes involved in the glycolysis process was found in the transcriptome and exhibited high expression levels (Fig. 4). The gene expression of pyruvate decarboxylase was up-regulated considerably, and it increased from 11.5 transcripts to 939.8 transcripts per million reads. This indicated that the anaerobic carbon metabolism in strain LZ-M might be achieved by glycolysis and pyruvate decarboxylation.

**Fig. 4.**
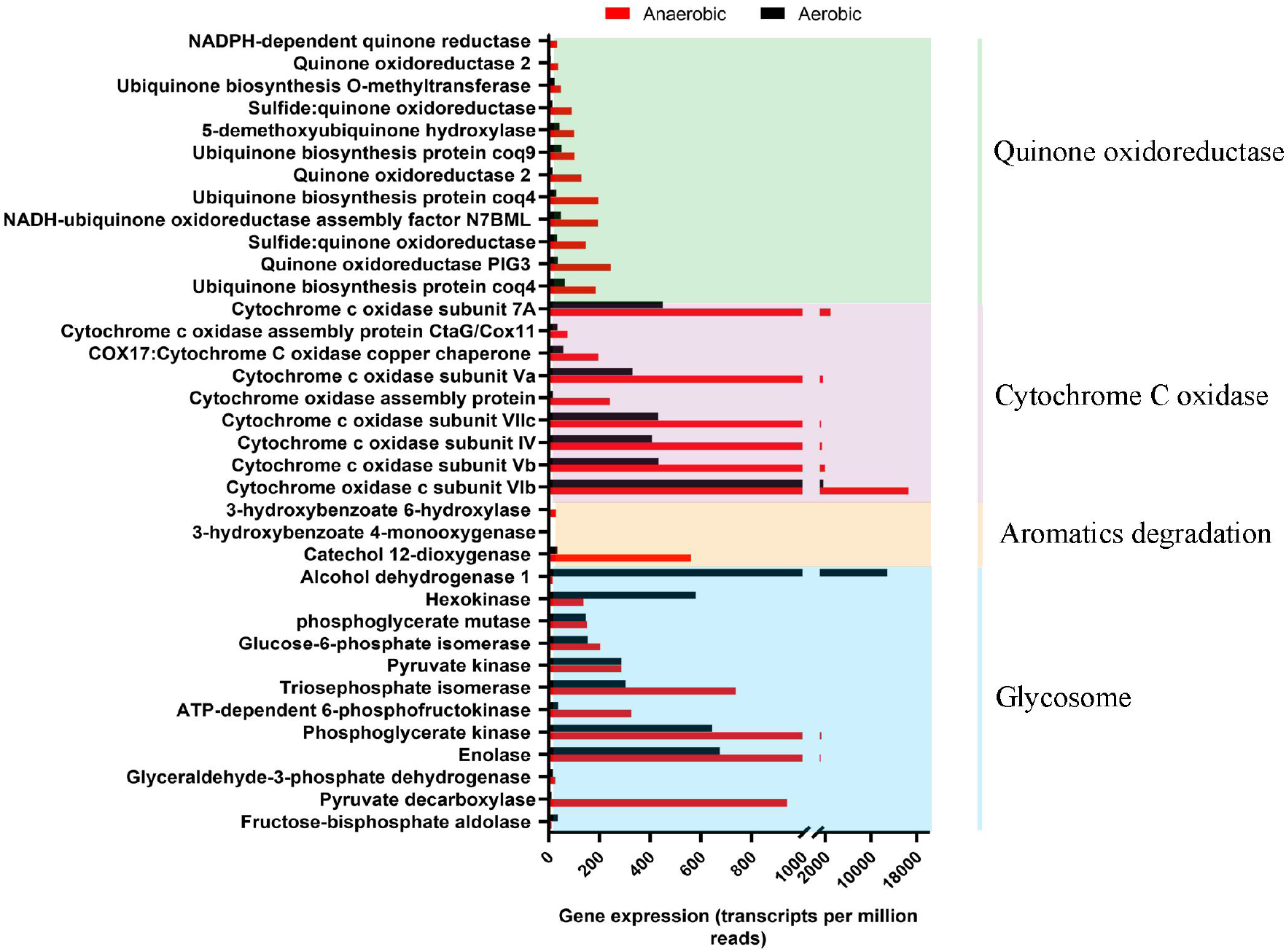
Expression counts of gene in *A. tabacinus* LZ-M after anaerobic and aerobic culture by transcriptome sequencing, including glycolysis process, aromatics degradation, cytochrome C oxidase and quinone oxidoreductase related genes.

In the degradation of AR73, the gene expression of enzymes involved in aromatics biodegradation, glycolysis and electron transfer were discovered, while the expression of gene involved in anaerobic decoloriztion or azoreduction was not found. The enzymes of aromatics biodegradation including 3-hydroxybenzoate 4-monooxygenase, 3-hydroxybenzoate 6-hydroxylase, and catechol 1,2-dioxygenase. Catechol 1,2-dioxygenase is able to degrade benzene, toluene and ethylbenzene (*37*). 3-hydroxybenzoate 4-monooxygenase and 3-hydroxybenzoate 6-hydroxylase are involved in benzoate degradation (*38, 39*). They may be involved in the metabolism of aniline and hydroxynaphthalene. Both aniline and naphthalene degradation intermediates salicylic acid, can enter the glycolysis process through glycosylation and phosphorylation for further metabolism (*40, 41*). In the glycolysis process, the splitting of the six-carbon glucose molecule into two pyruvate molecules by anaerobic oxidation (*42*). Pyruvate can be decarboxylated to acetaldehyde and CO_2_ by pyruvate decarboxylase in a stage of fungal anaerobic fermentation (*19*). Therefore, the CO_2_ generated in the anaerobic culture was considered as the result of anaerobic oxidation by glycolysis and fermentation (Fig. 3D). These results showed that strain LZ-M can completely mineralize AR73 to CO_2_ under anaerobic conditions.

During the glycolysis, NADH is produced and it need to regenerate NAD+ by dehydrogenation (*42*). NADH dehydrogenation requires electron transfer to the oxidized substrate, while the alcohol dehydrogenase that reduces acetaldehyde with NADH was rarely expression under anaerobic conditions (Fig. 4). Therefore, the regeneration of NAD+ may be combined with the cleavage of -C-N= bond of AR73.

Cytochrome C oxidoreductase and quinone oxidoreductases often act as mediators of electron transfer in cellular activities (*43*). Gene involved in the electron transfer reactions are found to increasing expression during anaerobic organic degradation (*22*). The up-regulation of cytochrome C oxidoreductase and quinone oxidoreductase may assist in electron transferring and electron balance during self-redox degradation of AR73. The self-redox reaction was represented as follow.

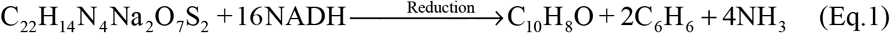

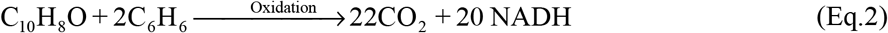

The degradation of AR73 was mainly achieved through carbon oxidation and nitrogen reduction. During the reduction process, 12 NADH are needed to decompose one molecule of AR73 to produce 4 molecules of NH_3_ (Eq.1). Similarly, hydroxynaphthalene and benzene analogs produced by one molecule of AR73 were completely mineralized to produce 22 molecules of CO_2_ and 20 NADH in oxidation process (Eq.2). It can be calculated by electron balance that when the self-redox equilibrium occurs, 80% of the carbon can be mineralized and converted into CO_2_. Azoreductase from bacteria and lignin peroxidase, manganese peroxidase and laccase from the fungi are able to degrade azo dyes (*33*), while they are not expression in strain LZ-M under anaerobic condition. Genes annotated as unknown function accounted for 51.81% of up-regulated transcripts. Thus, there may be new decolorizing enzyme among unknown genes that worth to explore.

### 3.5 Cloning and expression of unknown genes

The up-regulation genes were further screened based on the criteria that transcript expression level more than 150 counts and expression up-regulated fold more than 4.0, and then 140 genes was obtained. Among them, 51 genes were classified as oxidoreductase in enzymatic classification, 60 genes belonged to other classifications, and 29 genes belonged to unknown genes (Fig. 5A). Based on expression similarity (*44*), unknown genes have a high probability of being redox genes. Two unknown genes, named ord95 and ord118, have been cloned into *E.coli* Rosetta (DE3) and expressed by induction. The two proteins were purified successfully and detected by SDS-PAGE method (Fig. 5B). Alignment in NCBI database, the Ord95 gene showed the highest sequence identity of 58.71% with aconitate hydratase from *A. tubingensis*. Ord118 showed 100% similarity to hypothetical protein and no similarity to other functional proteins. The NADH dehydrogenase activity assay showed that both of them had NADH dehydrogenase activity (Fig. 5C and D). When the concentration of protein was 0.28 mg/L, the absorbance of NADH at A340 nm reduced at rate of 0.001 /min and 0.00043 /min by Ord118 and Ord95, respectively. Protein Ord95 and Ord118 were also found to have NADH-DCIP reductase activity using Biyuntian NADH oxidase detection kit (Fig. S3). These results suggest that both Ord95 and Ord118 are redox type enzymes. The enzyme activity analysis of AR73 decolorization using NADH as electron donor was conducted, and Ord95 had anaerobic decolorization enzyme activity (Fig. 5E and F), and the optimal reaction conditions of the enzymatic anaerobic decolorization were 37°C and pH 7.0 (Fig. S4). It couldn’t decolorize under aerobic conditions. When the concentration of AR73 was 25 mg/L and the concentration of Ord95 protein was 0.28 mg/L, the decolorization rate of AR73 was 0.4696 mg/L/h. This result confirms that Ord95 is a decolorizing reductase, and it might be the enzyme involved in the anaerobic decolorizing of AR73.

**Fig. 5.**
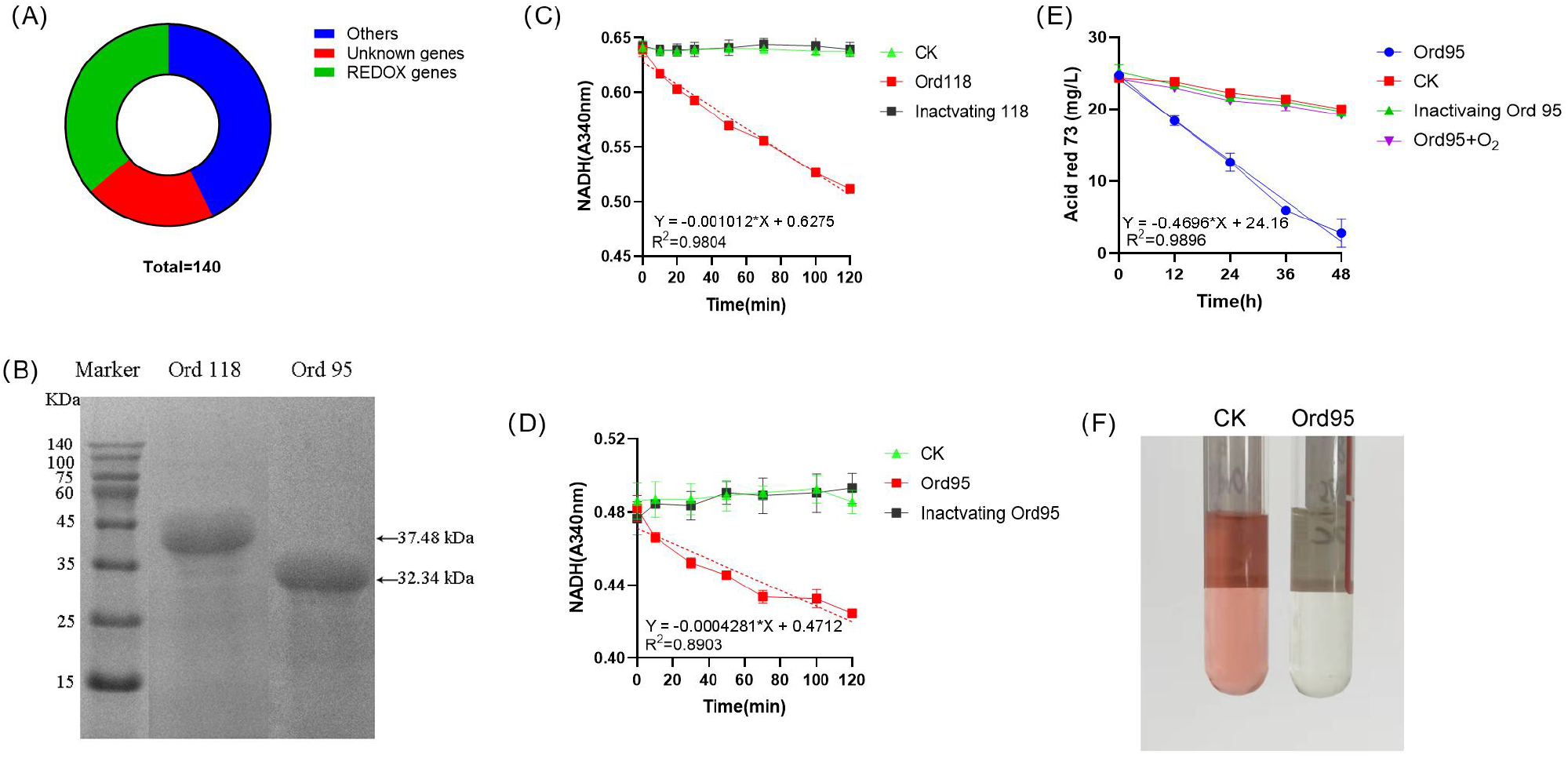
Unknown gene expression and function identification. (A) The distribution of oxidoreductase genes and unknown genes among the top 140 genes of up-regulated expression (count value > 150, up-regulated multiple > 4); (B) The unknown genes such as Ord95 and Ord118 after expression and purification were showed in the SDS-PAGE; (C, D) Dehydrogenase activity assay of Ord118 and Ord95 (mean ± SE, n = 3); (E-F) AR73 decolorization activity analysis of Ord95 (mean ± SE, n = 3). The concentrations of enzyme used were 0.28 mg/L.

### 3.6 Analysis of the decolorization mechanism of protein Ord95

In the degradation of AR73 by Ord95, aniline and phenylhydrazine were detected by HPLC in enzymatic reaction (Fig. 6A). The content of phenylhydrazine was increased at 24 h and decreased at 48 h, while the content of aniline kept increasing (Fig. 6B). This suggestion that Ord95 is the enzyme that broke the AR73 of -C-N= bond and its products were similar to fungal. Alignment in NCBI database, the 26-80 position of Ord95 sequence is similar to the glutathione S-transferase N-terminal domain with E-value of 7.07e-03, suggesting that this domain may be its active region. The main active site of glutathione transferase usually contains arginine (R) and tyrosine (Y) (*45*). The arginine and tyrosine near the domain region on the Ord95 were mutated to alanine (A), including 7Y, 8Y, 38R, 54R, 55R, 67Y. The 6 mutant proteins were expressed and purified (Fig. 6C), and their decolorizing activity was analyzed (Fig. 6D). Compared with Ord95 protein, the enzyme activity of mutants 38R>A, 54R>A, 55R>A decreased significantly to 43.1%, 17.43% and 36.9%, respectively. The enzyme activity of mutants 7Y>A, 9Y>A, 67Y>A did not change significantly. It suggested that 38R, 54R, 55R may be the active site of Ord95. Searching for template of 3D structure model of Ord95 in SWISS-MODEL, *C. albicans* actin binding protein 6m4c.1.A exhibited the highest quaternary structure quality estimate (QSQE) of 0.26, and the sequence identify was 32.98%. It suggests that Ord95 is a novel protein and its function has not been analyzed. Thus, 6m4c.1.A was selected as the template for use in homology model constructions of Ord95 by SWISS-MODEL. The functional domain of this protein was simulated as shown in the Fig. 6E and F. The model was a dimer with each monomer consisting of 1 beta sheet and 3 alpha helices.

**Fig. 6.**
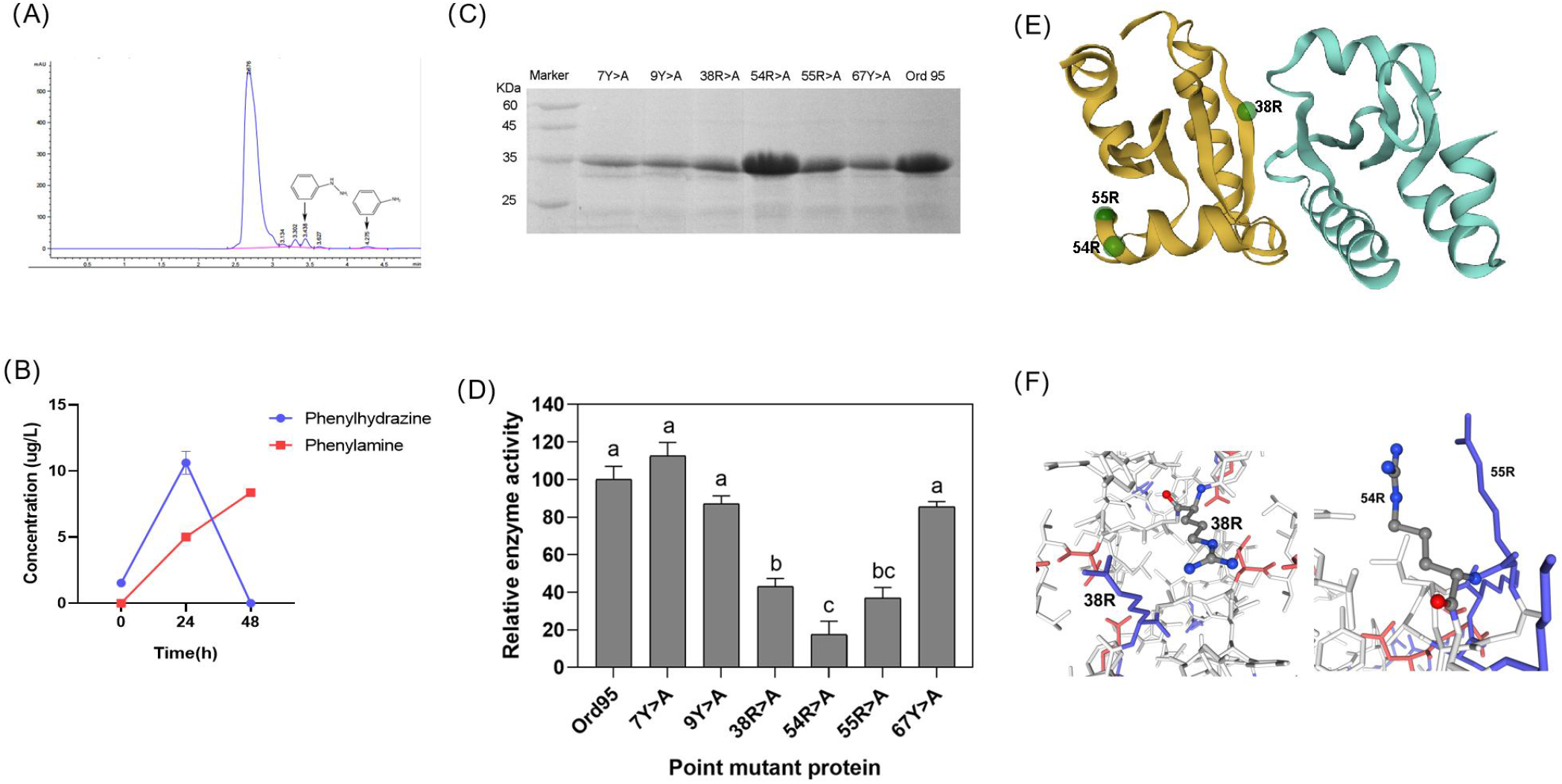
Products of enzymatic degradation of AR73 and structural prediction of Ord95 protein. (A) Determination of aniline and phenylhydrazine in AR73 enzymatic reaction solution by HPLC; (B) Changes of aniline and phenylhydrazine concentrations in AR73 enzymatic reaction solution with time (mean ± SE, n = 3); (C) SDS-PAGE image of Ord95 point mutant protein; (D) Relative enzyme activity of Ord95 point mutant protein (mean ± SE, n = 3); (E, F) Model constructions of Ord95 protein with SWISS-MODEL. The green dots are 38R, 54R, 55R mutation sites.

Three arginines were identified as the key sites of Ord95. The active site of glutathione transferases containing arginine is found in previous study (*45*). Arginine is an amino acid containing a guanidine group, which catalyzes the cycle of ornithine, promotes the formation of urea in organisms and turns ammonia into urea (*46*). Ord95 can aminate AR73 to form phenylhydrazine and aniline, which is associated with nitrogen metabolism. Therefore, the presence of arginine might promote the amination of nitrogen during AR73 degradation by Ord95. The degradation products of AR73 by the enzyme are similar to the degradation products by strain LZ-M, indicating that Ord95 is the functional enzyme for anaerobic decolorization. The metabolic mechanism of AR73 in strain LZ-M was represented in Fig. S5.

Accordingly, it was proved that strain LZ-M had a strong ability to completely mineralize azo dyes under anaerobic conditions, making up for the shortcomings of bacteria. This is the first report of the anaerobic degradation of azo dyes by obligately aerobic fungus. Besides, through the excavation of functional enzymes, new decolorizing enzymes were found, and the degradation mechanism of self-redox was revealed, which has not been reported in previous studies. Under anaerobic conditions, this self-redox degradation is rapid and complete. AR73 act as carbon and nitrogen source, reduced using NADH as an electronic donor and oxidized in glycolysis by strain LZ-M. Ord95 could cleave -C-N= in AR73 in the first step, and NADH generated during glycolysis can be delivered to Ord95. Fungi possess strong abilities to degrade refractory organic matter and secrete many function enzymes (*47, 48*). There are 128 *Aspergillus* genomes in the NCBI genome database, and 10,000-13,000 proteins were detected in each genome. Therefore, the excavation of fungal functional proteins has great application potential. However, the studies committed to organics degradation mechanism and functional proteins in fungi are still limited, and transcriptome analysis in this study revealed that a large number of proteins in strain LZ-M have not been identified yet. Furthermore, expanding of the protein library of fungi and the excavating functional enzymes might be focused in the further research.

## Acknowledgments

This work was supported by National Natural Science Foundation Grant (No: 31870082, 32070117), International Science and technology Cooperation project of Gansu Province (No: 2021-0204-GHC-0019), Gansu Science and Technology Association project (No: GKX20210506-16-5) and Lanzhou science and technology plan project (grant numbers: 2019-4-40). We thanks the Central Lab of School of Life Sciences, Lanzhou University for providing necessary equipment.

## Date availability

Transcriptome data of *A. tabacinus* LZ-M under aerobic culture and anaerobic culture is deposited at the Sequence Read Archive (https://www.ncbi.nlm.nih.gov/) under the accession numbers SRR14455363 and SRR14455360, respectively. The ITS gene sequence of *A. tabacinus* LZ-M is deposited under the accession numbers MZ127527 in NCBI (https://www.ncbi.nlm.nih.gov/search/all/?term=MZ127527). Data of LC-MS metabolite analysis are in the attachments named LC-MS_CK_vs_AR73_neg_info and LC-MS_CK_vs_AR73_pos_info.

